# Dominant-negative mutations in *CBX1* cause a neurodevelopmental disorder

**DOI:** 10.1101/2020.09.29.319228

**Authors:** Yukiko Kuroda, Aiko Iwata-Otsubo, Kerith-Rae Dias, Suzanna E.L. Temple, Koji Nagao, Lachlan De Hayr, Ying Zhu, Shin-Ya Isobe, Gohei Nishibuchi, Sarah K Fiordaliso, Yuki Fujita, Alyssa L. Rippert, Samuel W Baker, Marco L. Leung, Daniel C. Koboldt, Adele Harman, Beth A. Keena, Izumi Kazama, Gopinath Musuwadi Subramanian, Kandamurugu Manickam, Betsy Schmalz, Maeson Latsko, Elaine H Zackai, Matt Edwards, Carey-Anne Evans, Matthew C. Dulik, Michael F. Buckley, Toshihide Yamashita, W. Timothy O’Brien, Robert J. Harvey, Chikashi Obuse, Tony Roscioli, Kosuke Izumi

## Abstract

**Purpose:** This study aimed to establish variants in *CBX1*, encoding heterochromatin protein 1β (HP1β), as a cause of a novel syndromic neurodevelopmental disorder.

**Methods:** Patients with *CBX1* variants were identified, and clinician researchers were connected using GeneMatcher and physician referrals. Clinical histories were collected from each patient. To investigate the pathogenicity of identified variants, we performed *in vitro* cellular assays, neurobehavioral and cytological analyses of neuronal cells obtained from newly generated *Cbx1* mutant mouse lines.

**Results:** In three unrelated individuals with developmental delay, hypotonia, and autistic features, we identified heterozygous *de novo* variants in *CBX1*. The identified variants were in the chromodomain, the functional domain of HP1 β, which mediates interactions with chromatin. *Cbx1* chromodomain mutant mice displayed increased latency-to-peak response, suggesting the possibility of synaptic delay or myelination deficits. Cytological and chromatin immunoprecipitation experiments confirmed the reduction of mutant HP1β binding to heterochromatin, while HP1β interactome analysis demonstrated that the majority of HP1β-interacting proteins remained unchanged between the wild-type and mutant HP1β.

**Conclusion:** These collective findings confirm the role of *CBX1* in developmental disabilities through the disruption of HP1β chromatin binding during neurocognitive development. As HP1β forms homodimers and heterodimers, mutant HP1β likely sequesters wild-type HP1β and other HP1 proteins, exerting dominant-negative effects.

## Introduction

The heterochromatin protein (HP1) family are evolutionarily conserved structural proteins that regulate the formation of heterochromatin, an inactive genomic region.^1^ In humans, three highly homologous HP1 proteins (HP1α, HP1β, and HP1γ) have functionally redundant and non-redundant roles.^2^ HP1 proteins form homo- and heterodimers, which bind to methylated histone 3 lysine 9 residues (H3K9) and bring two methylated H3K9s into close proximity, resulting in chromatin compaction.^3^ In addition, HP1 proteins interact with various chromatin regulatory molecules, including euchromatic histone lysine methyltransferase 1 (EHMT1), chromodomain helicase DNA binding protein 4 (CHD4), and activity-dependent neuroprotector protein (ADNP), and cooperatively regulate chromatin organization.^4,5^

Precise regulation of heterochromatin formation is an essential genome regulatory mechanism and disruption of this process is increasingly recognized as a cause of neurodevelopmental disorders. Among these, genetic variants in HP1-interacting proteins represent an important class of neurodevelopmental disorders, including Kleefstra syndrome due to *EHMT1* mutations,^6,7^ Sifrim-Hitz-Weiss syndrome due to *CHD4* mutations,^8^ Helsmoortel-van der Aa syndrome due to *ADNP* mutations,^9^ Weiss-Kruszka syndrome due to *ZNF462* mutations,^10^ and Luo-Schoch-Yamamoto syndrome due to *RNF2* mutations.^11^ All of these syndromes are associated with developmental delay and intellectual disability, suggesting the importance of HP1-interacting proteins in normal neurocognitive development. Although HP1β participates in the regulation of brain development,^12^ dysfunction of the HP1 protein itself has not until now been associated with human neurocognitive disorders.

Here, we report three individuals with developmental disabilities due to *de novo* missense variants in *CBX1*, which encodes HP1β. The identified variants were found to alter the intracellular localization of HP1β and abolished the interaction between HP1β and histone-repressive marks, confirming the importance of *CBX1* and precise HP1β-heterochromatin interactions in human neurocognitive development.

## Material and Methods

### Cohort Recruitment

Through matchmaking via GeneMatcher^13^ and international collaborative efforts, a cohort of affected individuals with *CBX1* variants and an overlapping phenotype was assembled. These individuals were referred from different collaborating institutes in the USA and Australia. Detailed clinical information was ascertained through recontact or detailed review of medical records by neurologists, pediatricians, clinical geneticists and genetic counsellors. Exome sequencing methods are described in the Supplementary Method.

### *CBX1* variant molecular modelling

Molecular modelling was completed using the structure of the HP1β chromodomain bound to a histone H3K9Me3 peptide (PDB 6D07^14^). The ChimeraX molecular graphics system^15^ was used to visualize the structure and describe the intra- and inter-molecular forces. Modelling of the W52L, N57D and T51P substitutions was accomplished using the Swapaa command using the Dunbrack backbone-dependent rotamer library^16^ and taking into account the lowest clash score, highest number of H-bonds and highest rotamer probability. ChimeraX functions were used to display all hydrogen and π bonds, contacts, clashes and electrostatic properties of the protein surface.

### Cell culture

Lymphoblastoid cell lines (LCLs) were obtained from individual 1 and individual 2. Gender-ethnicity matched control LCL samples were used as controls. LCLs were cultured in RPMI 1640 with 300mg/L L-glutamine (Life Technologies, 11875085) supplemented with 20% HyClone FBS (Fisher Scientific, SH3007103), 0.2% penicillin-streptomycin (Life Technologies, 15140122), 0.2% Plasmocin (Invivogen, ant-mpp), and 1% Glutamax (Life Technologies, 35050061). The HEK293T cell line was cultured in DMEM with 4.5 g/L D-Glucose and 110mg/L sodium pyruvate (Life Technologies, 11360070) supplemented with 10% HyClone FBS, and 0.2% penicillinstreptomycin. All cells were cultured at 37°C in 5% CO2. CBX1 expression vectors were transfected into HEK293T cells using Lipofectamine 2000 (Life Technologies, 11668-030) according to the manufacturer’s protocol. The Myc-DDK-tagged-CBX1 expression vector was purchased from Origene (RC205672). *CBX1* mutations were introduced using the Q5 Site-Directed Mutagenesis Kit (New England Biolabs Inc., E0554S) following the manufacturer’s protocol. Sanger sequencing confirmed the suspected change and ruled out additional, nonspecific changes. Immunoblotting and chromatin immunoprecipitation Western blot (ChIP-WB), and immunofluorescent (IF) staining methods are described in the Supplemental Method.

### Proteomics analyses

To establish stable cell lines for inducible expression of Flag-tagged HP1 β, the Flp-In T-REx 293 system (Invitrogen) was used as described.^17^ Protein extraction, immunoprecipitation, and mass spectrometry were performed as described.^17^ The details of the proteomics analysis methods are described in the Supplemental Method.

### Generation of *Cbx1* mutant mice by CRISPR/Cas9 genome editing

CRISPR/Cas9-mediated gene editing was conducted in zygotes of C57BL/6 mice. A guide RNA targeting a genomic region of interest, Cas9 and ssDNA containing mutant sequence were electroporated into zygotes. The details of genome editing, primary neuronal culture and Nissel staining methods are described in the Supplementary Method.

### Behavioral testing

Thirteen *Cbx1* W52L+/− and 10 +/+ male littermates were tested in a battery of behaviors, starting at approximately 4 months old. Procedures and timeline used to test *Cbx1* W52L+/− line as follows: open field on day 1, handling and Y-maze spontaneous alternations on days 2-5, accelerating Rotarod on days 7-9, acoustic startle (Pre-pulse Inhibition) on day 10. Detailed protocols are described in the Supplemental Method.

## Results

### Clinical description of individuals with *CBX1* variants

The clinical features of three patients were compared, and their variant status was summarized (**Table 1**). Shared clinical features included developmental delay, hypotonia, autistic features, and variable dysmorphic features. The three identified variants, p.N57D, p.W52L, and p.T51P, revealed an *in silico* score interpretation consistent with an increased chance of pathogenicity with VarCards combined scores of 19/23, 21/23, and 22/23, respectively.^18^ Additionally, these variants are not present within the gnomAD database.^19^ Thus, they are rare and likely deleterious variants.

**Table 1:** Clinical features of three individuals with *CBX1* variants.

|  | Individual 1 | Individual 2 | Individual 3 |
| --- | --- | --- | --- |
| Gender | Female | Male | Female |
| Age at initial genetics evaluation | 1 year 6 months | 5 years | 5 months |
| *CBX1* variant | c.169A>G, p.Asn57Asp<br>(NM_001127228.2) | c.155G>T, p.Trp52Leu<br>(NM_001127228.2) | c.151A>C, p.Thr51Pro<br>(NM_001127228.2) |
| Genome coordinate | chr17(hg19):g.46153512 T>C | chr17(hg19):g.46153526C>A | chr17(hg19):g.46153530T>G |
| Age at last visit | 6 years | 5 years | 7 years |
| OFC at last visit | 51.5 cm (80<sup>th</sup> percentile)(at 4 years 9 months) | 52.8 cm (85-97<sup>th</sup> percentile) | 52.0 cm (91<sup>st</sup> percentile) |
| height at last visit | 118.4 cm (85<sup>th</sup> percentile) | 103.7 cm (50-70<sup>th</sup> percentile) | 9<sup>th</sup> centile |
| weight at last visit | 33 kg (>99<sup>th</sup> percentile) | 17 kg (50-70<sup>th</sup> percentile) | 77<sup>th</sup> centile |
| Feeding difficulties | Present as neonate; attributed to ankyloglossia s/p frenotomy and reduced maternal milk supply | Colicky baby | Poor feeding as infant; overeating by 15 months (continues today) |
| Growth | Obesity (weight >99<sup>th</sup> percentile, BMI >99<sup>th</sup> percentile) | 50<sup>th</sup>-75<sup>th</sup> percentiles for height and weight | Elevated BMI given weight/length, but parents also control intake strictly |
| Overall Development | Global developmental delay | Global developmental delay | Global developmental delay |
| Motor abilities | Hypotonia, gross motor delay (sitting at 10m, pulling to stand at 14m, and walking at 23m); still clumsy | Cardiorespiratory arrest with RSV bronchiolitis aged 6 weeks thought to have caused developmental delay initially | Initial delays (walked at age 4) but significantly improved; now has good strength but mild hyperflexibility, indicating hypotonia |
| Cognitive abilities | Mild-to-moderate intellectual disability | Mild-to-borderline disability probable | Moderate-severe intellectual disability |
| Behavioral Issues | Autism spectrum disorder, obsessive-compulsive behaviors | Autism spectrum disorder | Autism spectrum disorder (more severe range) |
| Verbal abilities | Expressive language delay, non-verbal, uses AAC to communicate | Expressive language delay | Non-verbal; occasionally use AAC to communicate, but only recently |
| MRI | Normal brain MRI at 3y | Enlarged extra-axial fluid spaces | Normal (age 3) |
| Muscle tone | Hypotonia | Hypermobility and hypotonia | Good strength but hypermobile |
| Ear Problems | Recurrent/chronic otitis media s/p<br>bilateral myringotomy tubes | None reported | None reported |
| Craniofacial<br>Abnormalities | Frontal bossing, midface<br>hypoplasia, round-shaped face,<br>epicanthal folds, upslanting<br>palpebral fissures, flat nasal bridge<br>with short nose, tented upper lip<br>with thinning of the lateral upper lip,<br>closely spaced nipples, short fingers<br>and toes, 2-3 syndactyly of toes<br>bilaterally | Arched eyebrows, broad forehead,<br>tall skull with flat occiput, small<br>overfolded ears. Increased extra-axial CSF and mild ventricular<br>dilatation | Frontal bossing, round face,<br>almond-shaped eyes |
| Genitourinary<br>Issues | No issues | Incomplete testicular descent<br>treated by orchiopexy | None reported |
| Skeletal<br>Issues | No issues | Narrow flat feet. Right hip pain for<br>months led to MRI showing a right<br>intramedullary femur lesion. Biopsy<br>showed inflammation. No evidence<br>of bacterial infection and<br>subsequent slow progression<br>suggests chronic relapsing<br>multifocal osteomyelitis | None reported |
| Gastrointestinal Issues | None reported | None reported | Significant GI bloating/distension;<br>improved with celiac-free diet |

### HP1β protein modelling

In order to understand the functional impact of the p.W52L, p.N57D and p.T51P substitutions, we utilized molecular modelling using the structure of the HP1β chromodomain bound to a histone H3K9Mme3 peptide (PDB 6D07).^14^ Residue W52 occurs in a β-sheet region that plays a vital role in stabilizing the organization of three parallel sheets (**Fig 1a**). W52 forms a cation-π interaction with the R30 residue and has numerous contacts with other residues in the parallel β-sheets, including D28 and L39. The p.W52L substitution disrupts both the cation-π interaction with R30 and contacts with the side chains of D28 and L39 (**Fig 1b**) as well as creates clashes with R30. Hence, the p.W52L substitution is predicted to disrupt the β-sheet structure in a region that forms the binding interface for the methylated histone mark. However, p.W52L does not directly alter the structure of the key H3K9Me3-binding aromatic cage. In contrast, the N57 residue lies directly within the HP1β histone binding cleft. The N57 side chain forms a hydrogen bond with the backbone of the histone at R8, as well as HP1β E53, which forms a hydrogen bond with histone S10 (**Fig 1c**). The p.N57D substitution results in the exchange of a positively-charged nitrogen for a negatively-charged oxygen in the side chain. Conformational changes between N57 and D57 are predicted to cause the loss of a hydrogen bond between D57 and the backbone of the histone peptide, and the loss of a second hydrogen bond with CBX1 E53 (**Fig 1d**). The lost hydrogen bond with E53 is replaced by a new hydrogen bond with histone S10. We also examined the charge state of the HP1β histone-binding pocket (**Fig 1e and 1f**). This appears to be close to neutral due to oxygen and nitrogen atoms in N57, while the surrounding area has a greater negative charge (**Fig 1e**). However, N57D results in two oxygen atoms in the aspartic acid residue, which acts to increase the negative charge of the histone binding cleft (**Fig 1f**), which may impact stability of the HP1 β-histone interaction.^20,21^ Lastly, residue T51 is located at the beginning of the final beta-sheet in the chromodomain, where it forms contacts with neighboring residues, including the histone binding residues W42 and E53 (**Fig 1g**). The p.T51P substitution is predicted to be detrimental to the structure of this region due to the loss of stabilizing H-bonds with E53 and W42, although the substitution forms numerous new contacts with W42. In addition, the proline ring imposes constraints on the beta-strand formation by using one available carbon donor and fixing the bond angles at the carbon backbone (**Fig 1h)**.^22^

**Fig 1:**
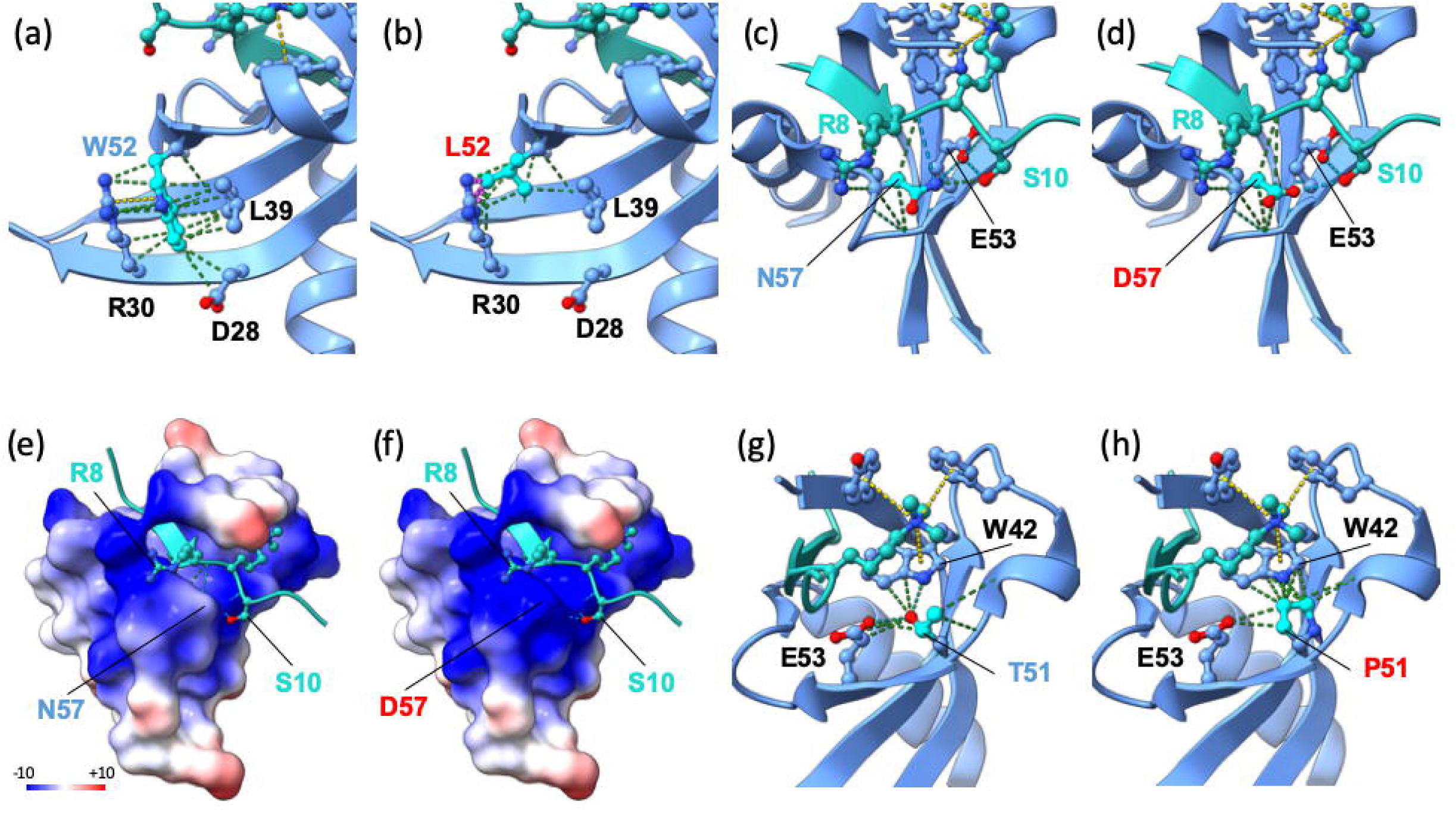
Molecular modelling of *CBX1* missense variants. (a)(b) W52 forms a number of interactions with nearby residues (D28, L39) in addition to a cation-π interaction with R30 that spans across three proximal β-sheets. Substitution W52L removes interactions with the side chains of D28 and L39 and creates clashes with R30. (c)(d) Substitution N57D results in the loss of a hydrogen bond between D57 and the backbone of the histone peptide, and the loss of a second hydrogen bond with HP1β E53. (e)(f) Substitution N57D also increases the negative charge on the surface of the histone-binding pocket. Charge is indicated as blue for negative and red for positive charge ranging from −10 to +10 kcal/(mol·e). (g)(h) Substitution T51P results in the loss of stabilizing H-bonds with E53 and increases unfavorable contacts with W42.

### *CBX1* variants impact HP1β-heterochromatin interaction

Molecular modeling is consistent with the identified *CBX1* variants abolishing the interaction between HP1β and methylated histones. Within the nucleus, wild-type (WT) HP1β colocalizes with 4’,6-diamidino-2-phenylindole (DAPI) dense regions known as chromocenters. These regions are composed of constitutive heterochromatin.^23^ Previous studies have demonstrated that HP1β protein lacking the CD or harboring a deleterious variant within the CD (HP1β V23M) localizes throughout the nucleus instead of accumulating in DAPI dense regions.^23,24^ Thus, we tested the nuclear localization of HP1β with patient-identified variants to determine if it localizes throughout the nucleus in a manner similar to the previously reported CD mutant HP1β. Using FLAG-CBX1 cDNA overexpression in HEK293T cells, we evaluated the intranuclear distribution of WT and mutant HP1β by immunofluorescence using an antibody against FLAG. While WT HP1β colocalized with DAPI dense regions, signals of HP1β with CD variants (T51P, W52L, and N57D) showed homogeneous nuclear staining, indicating mislocalization of the mutant HP1β, similar to that observed for the artificial V23M mutant (**Fig 2a**).

**Fig 2:**
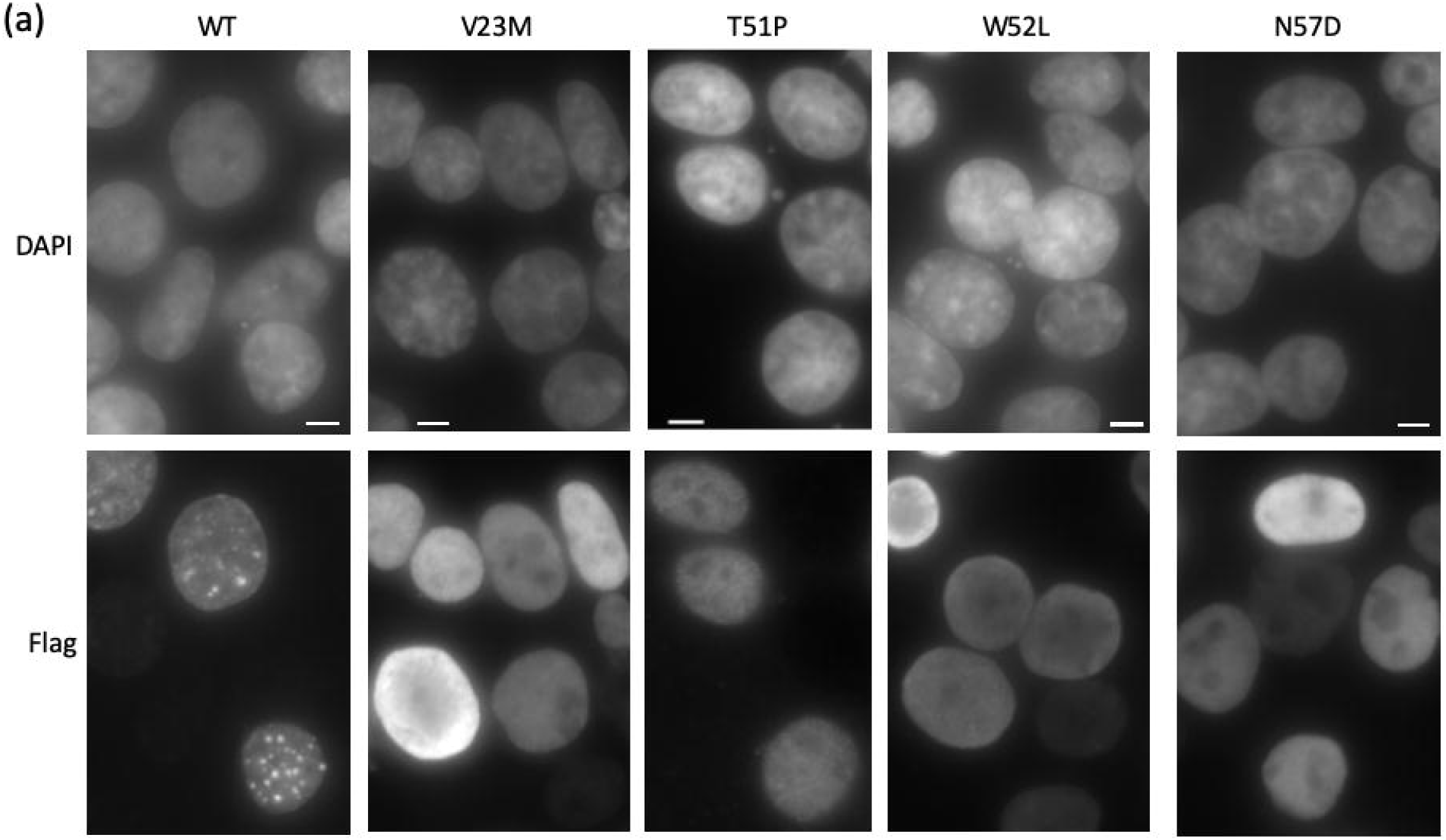
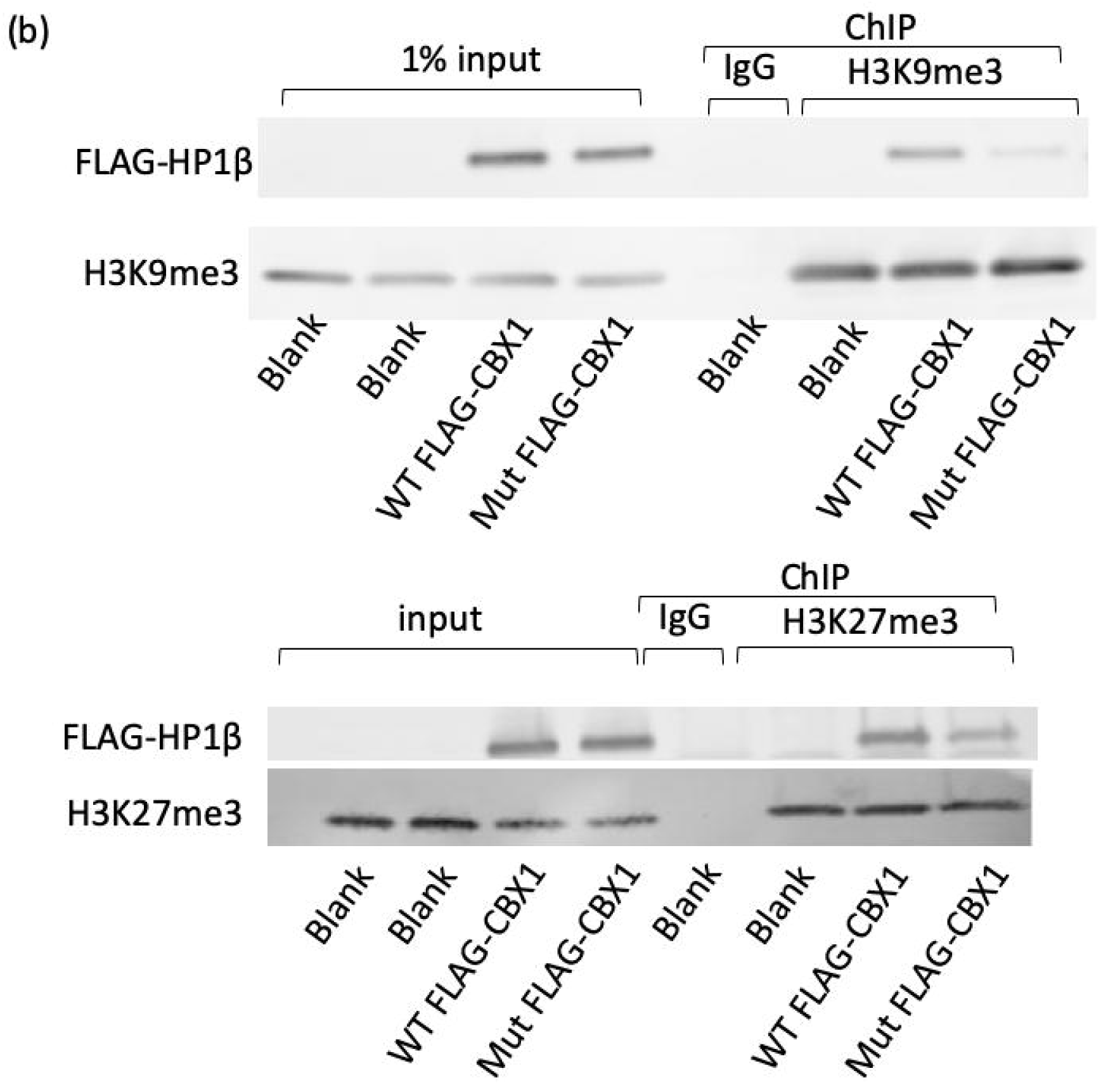
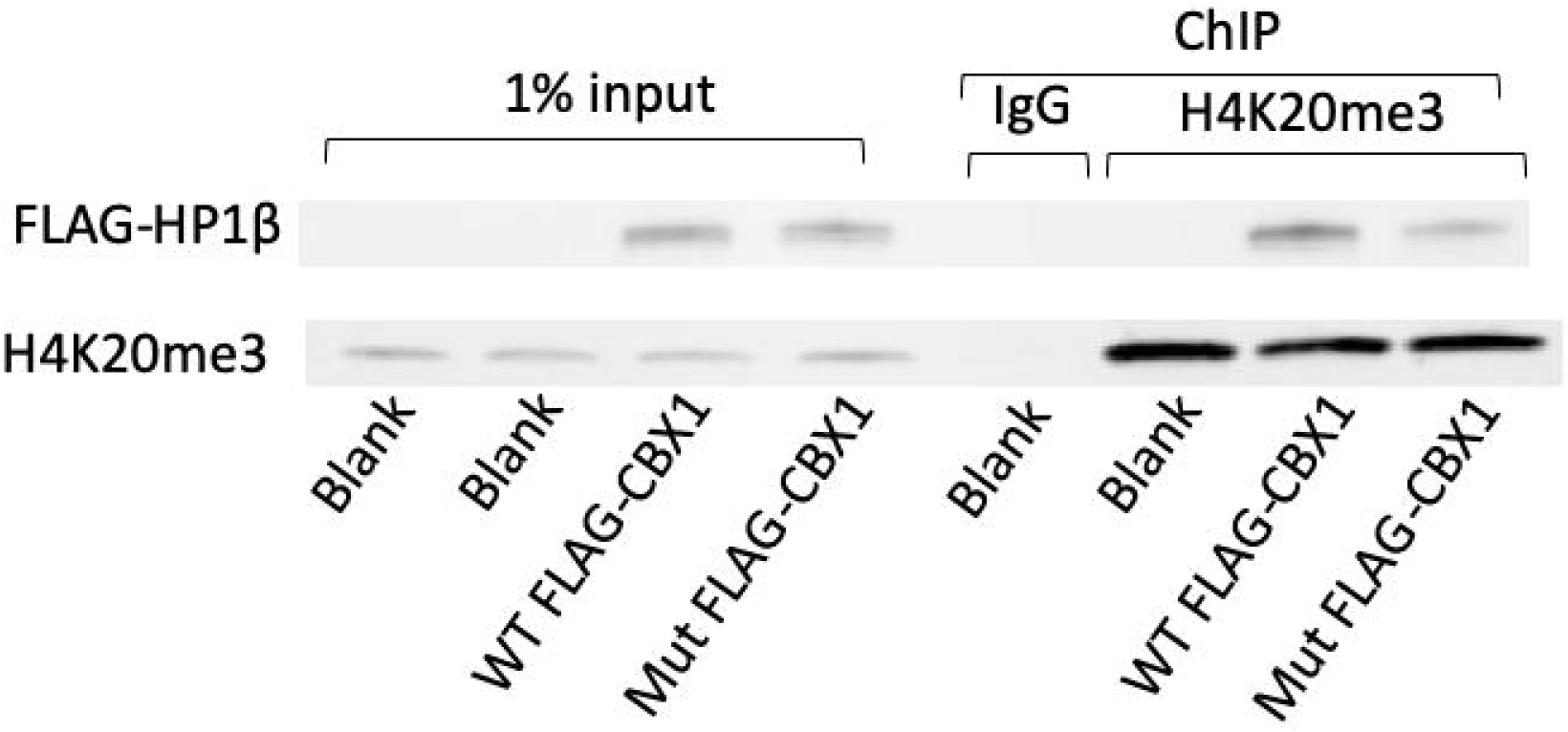
Mutant HP1β abolishes HP1β-chromatin interactions. (a) Mislocalization of mutant HP1β. CD mutant HP1β demonstrates diffuse HP1β signal within nucleus, although WT HP1β signals co-localize with DAPI-dense regions. In all the immunofluorescence experiments, at least two biological duplicates were performed. Scale bar is 5μm. (b) Reduced HP1β binding to repressive chromatin marked histones. Reduced H3K9me3, H4K20me3 and H3K27me3 binding of mutant HP1 β. ChIP was performed after 48 hours of FLAG-*CBX1* cDNA overexpression (wildtype and N57D mutant). Although equal amount of FLAG-HP1β is expressed between wild-type and N57D mutant, less N57D mutant was identified in H3K9me3/H4K20me3/H3K27me3 marked chromatin fraction compared to control. Biological duplicates revealed the consistent result. Similar results were obtained with the W52L mutant for H3K9me3 (**Supplementary Fig 1**).

As DAPI dense regions represent foci of heterochromatin, it was hypothesized that mutant HP1β binds less efficiently to compacted heterochromatin. Using WT and N57D mutant (Mut) FLAG-CBX1 cDNA vectors, the amount of H3K9me3-bound HP1β was evaluated by chromatin immunoprecipitation (ChIP)-Western blot.^25^ This analysis showed a reduction in Mut FLAG-CBX1 binding to H3K9me3 relative to WT (**Fig 2b**). Similar results were observed for H4K20me3 and H3K27me3, which are histone modifications seen in facultative heterochromatin (**Fig 2b**). Similar results were also obtained for the W52L mutant for H3K9me3 chromatin binding (**Supplementary Fig 1**). These observations are consistent with the notion that the CD mutants reduce the ability of HP1β to bind methylated histones found in heterochromatin.

### Impacts of *CBX1* variants in patient-derived lymphoblastoid cells

The effects of *CBX1* variants were also evaluated in patient-derived lymphoblastoid cell lines (LCLs). LCLs were generated from blood samples obtained from individuals 1 (N57D) and 2 (W52L). Immunoblotting showed that HP1β levels were increased for these HP1β variants (**Supplement Fig 2a**). HP1β immunofluorescence staining of these samples was performed. There were no discrete DAPI dense regions observed in these LCLs and the intracellular distribution pattern did not appear to be different between patient and control samples (**Supplement Fig 2b**).

### Proteomic analyses of mutant HP1β

HP1 interacts with methylated lysine of histones using the CD, whereas the C-terminal chromo shadow-domain (CSD) interacts with other HP1 proteins.^26,27^ As differences were observed in the intracellular accumulation of HP1β proteins and intranuclear distribution due to the *CBX1* mutations, the impact of HP1β CD mutations on protein-interacting partners were investigated by immunoprecipitation followed by mass spectrometry (IP-MS). This analysis was performed using T-REx 293 cells stably expressing FLAG-WT CBX1, or FLAG-mut CBX1 (W52L or N57D). It has previously been established that a CD mutant HP1 β (V23M) and a CSD domain mutant (I161E) abolish HP1 dimer formation.^28^

The effect of the HP1β N57D and W52L mutations on the ability to form HP1 hetero-or homodimers was investigated. WT and both CD mutants (N57D and W52L) were able to interact with HP1α and HP1γ, whereas the CSD mutant (I161E) failed to interact with them (**Fig 3**). While immunoprecipitation of the I161E mutant demonstrated that the majority of the identified proteins were HP1β, the identified HP1 most likely represented the immunoprecipitated FLAG-HP1β itself, rather than I161E preserving its ability to form an HP1β homodimer. Comparison of the total amount of immunoprecipitated HP1 proteins, including HP1α, HP1β, and HP1γ, demonstrated that I161E was associated with a reduced amount of total HP1 protein compared to the WT and the HP1 β CD mutant, although the amount of immunoprecipitated HP1 β was comparable among WT, CD mutant, and CSD mutant HP1β (**Fig 3b**).

**Fig 3:**
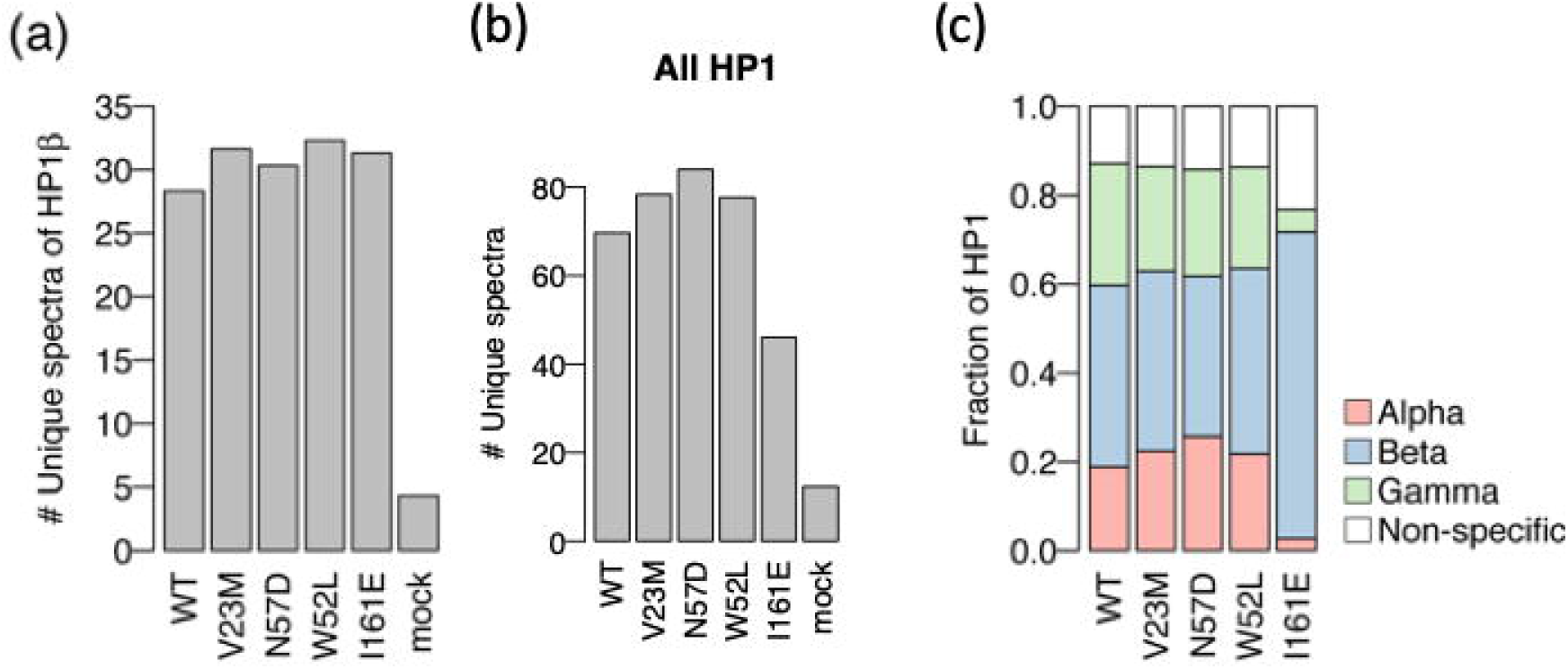
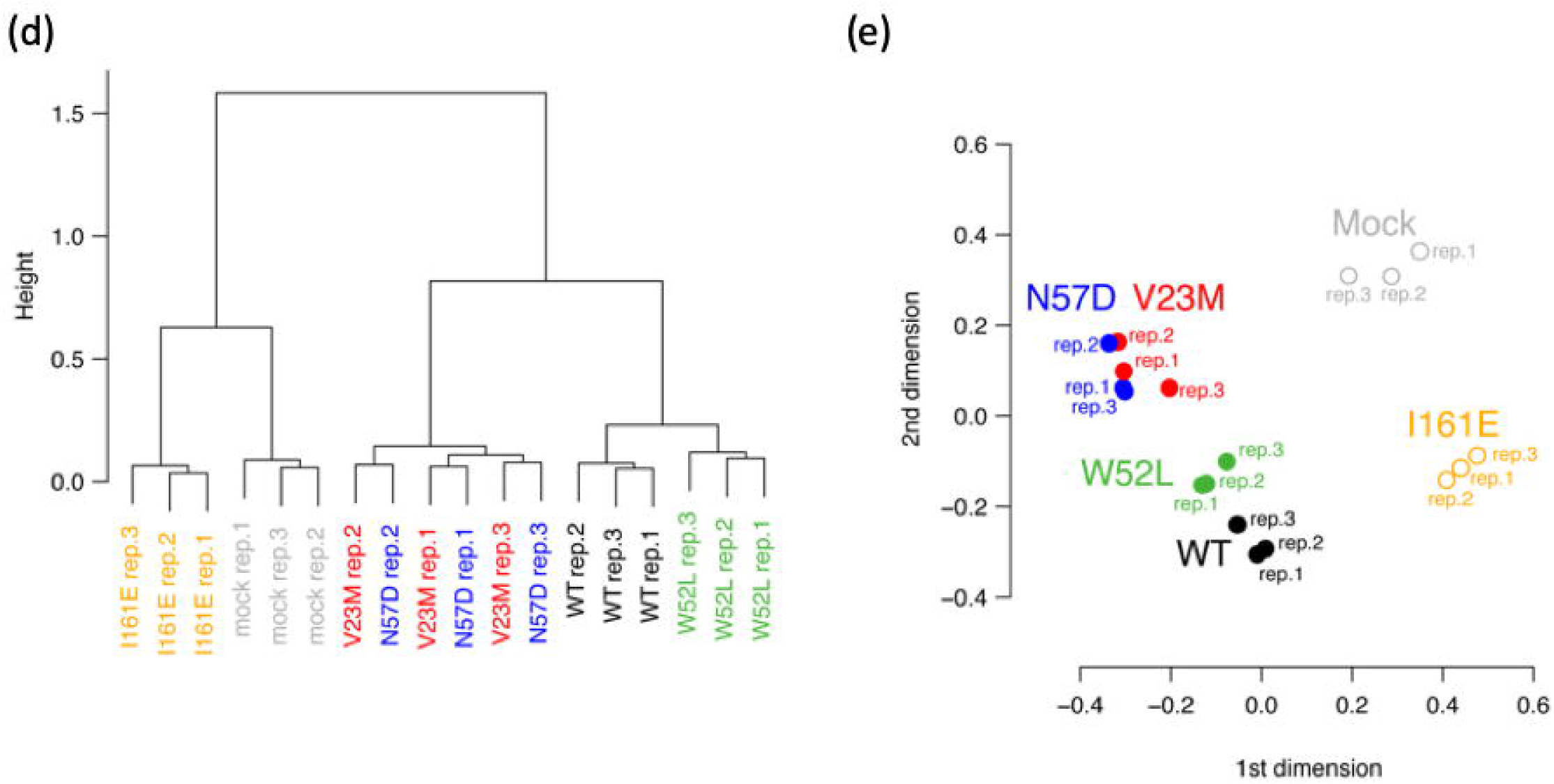
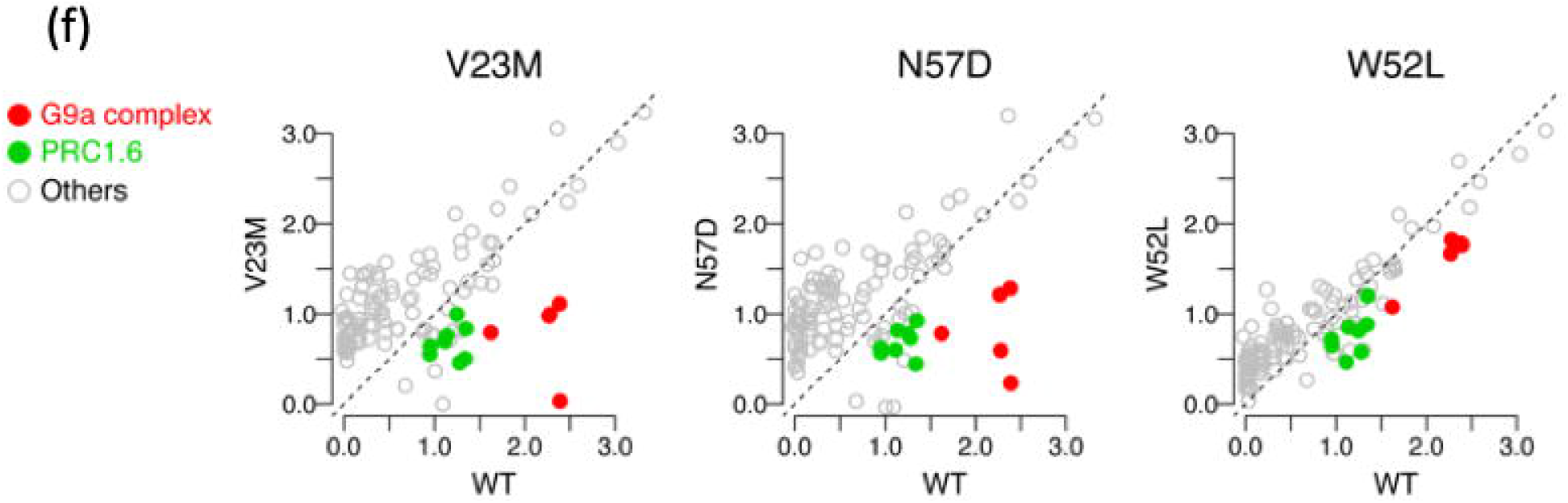
Evaluation of HP1β-interacting proteins using IP-MS. The abundance of HP1β interacting molecules were compared for wild-type (WT), V23M, W52L, N57D and I161E HP1β. (a) Confirmation of successful FLAG-HP1β immunoprecipitation (IP). Average number of unique spectra specific to HP1β was comparable among WT and mutant HP1β. (b) Total number of unique spectra specific to HP1. Note the reduction of HP1 count in the CSD mutant (I161E) HP1β, suggesting a lack of ability to form homo and hetero HP1 dimers. (c) Fraction of HP1 protein bound to FLAG-HP1β. While CSD mutant HP1β (I161E) abolished its binding to HP1α and HP1γ, CD mutant HP1β (V23M, W52L, and N57D) preserved the ability to bind to HP1α and HP1γ. The average of three IP-MS replicates is shown. (d) Hierarchical clustering analysis of FLAG-HP1β interacting protein profile revealed that N57D mutant had a similar profile to V23M, while W52L demonstrated a similar profile to WT HP1β. Mock represents negative control using parental cells not expressing Flag-HP1β. (e) Multi-dimensional scaling analysis of FLAG-HP1β interacting protein profile revealed that W52L sample shows intermediate phenotype between WT and other CD HP1β mutant samples (V23M and N57D). (f) Comparison of IP enrichment for 142 HP1β-interacting proteins between mutant HP1β to the WT control. V23M and N57D mutant samples revealed a lack binding to the G9a complex (red) and PRC1.6 (green). W52L samples demonstrated a mild reduction of binding to the G9a complex and PRC1.6. G9a complex: EHMT1, EHMT2, WIZ, ZNF462, ZNF644.^29–31^ PRC1.6: E2F6, L3MBTL2, MGA, PCGF6, RING1, RNF2, TFDP1, WDR5.^32^

Hierarchical clustering and multi-dimensional scaling analyses showed that the HP1β interacting molecule profiles were very similar between V23M and N57D (**Fig 3d and 3e**). In contrast, those of W52L were clustered differently relative to those of V23M and N57D. Multi-dimensional scaling analyses revealed that the interactome profile of W52L was intermediate between that of WT and other CD mutants (V23M and N57D) (**Fig 3d and 3e**).

Comparison of the list of HP1β interacting molecules between WT and CD mutant HP1β revealed that the V23M and N57D mutants demonstrated impaired binding to the GLP/G9a complex (EHMT1, EHMT2, WIZ, ZNF462, and ZNF644) ^29–31^ and the PRC1.6 complex (E2F6, L3MBTL2, MGA, PCGF6, RING1, RNF2, TFDP1, and WDR5) ^32^ (**Fig 3f**). In contrast, the abundance of other HP1 β interacting molecules was higher in the CD mutant than in the WT (**Fig 3f**). Consistent with the findings of multi-dimensional scaling analysis, the degree of W52L HP1 β binding changes was milder than those of V23M and N57D (**Fig 3f**).

### Mouse *Cbx1* W52L and N57D mutant phenotypes

The effects of *CBX1* variants *in vivo* were investigated by establishing *Cbx1* mutant mouse lines harboring patient-identified variants (W52L and N57D) using CRISPR/Cas9 genome editing. These codons are conserved between humans and mice (**Fig 4**). Heterozygous *Cbx1* mutant mice were born following the expected Mendelian ratio by crossing a heterozygous *Cbx1* mutant mouse and a WT mouse, although homozygous *Cbx1* mutant mice were not obtained from mating heterozygous W52L male and female mutant mice (at least 28 pups born from five litters, constituting 21 heterozygous (W52L +/− and 7 WT pups), suggesting that homozygous HP1β CD variants cause embryonic lethality.

**Fig 4:**
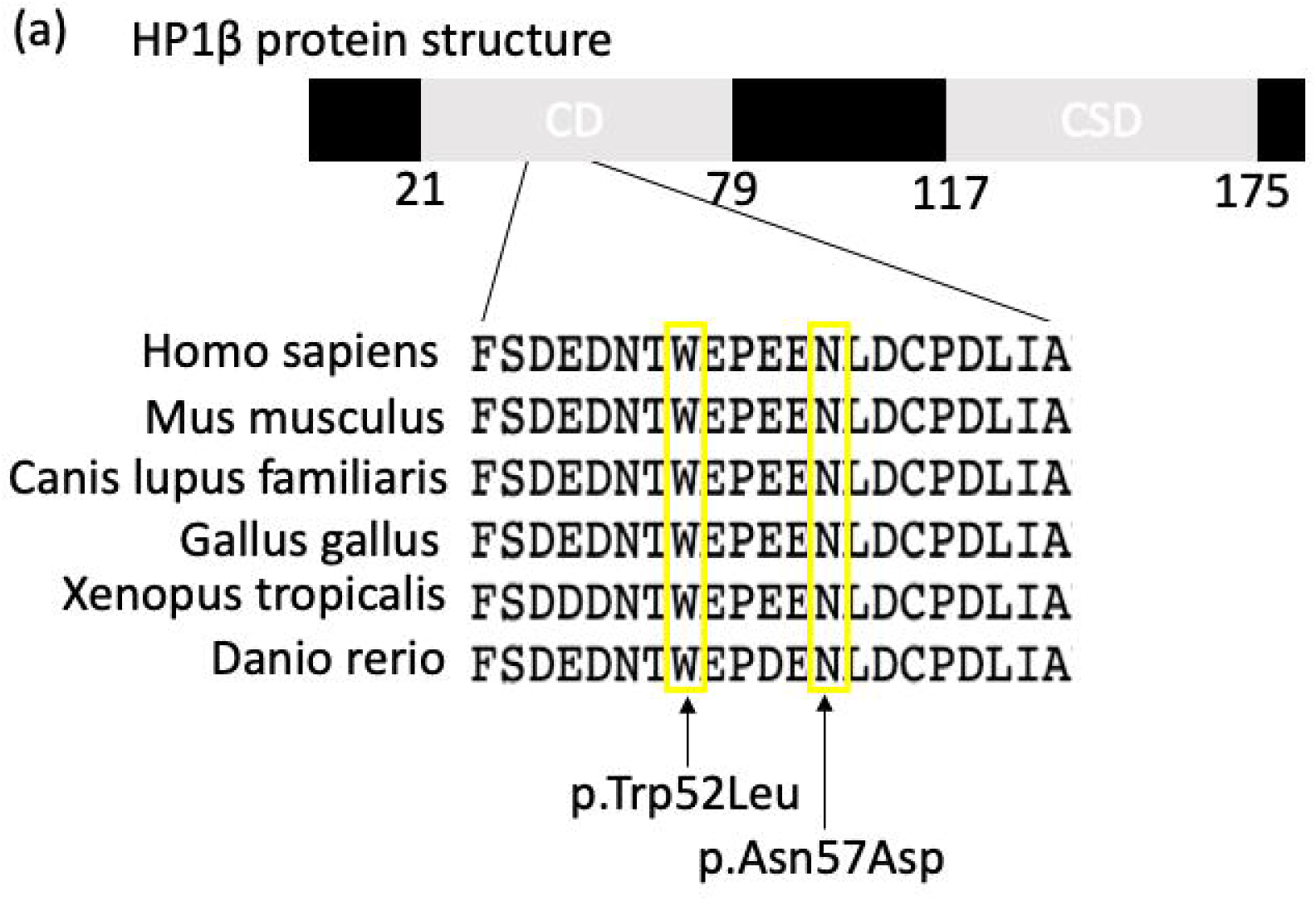
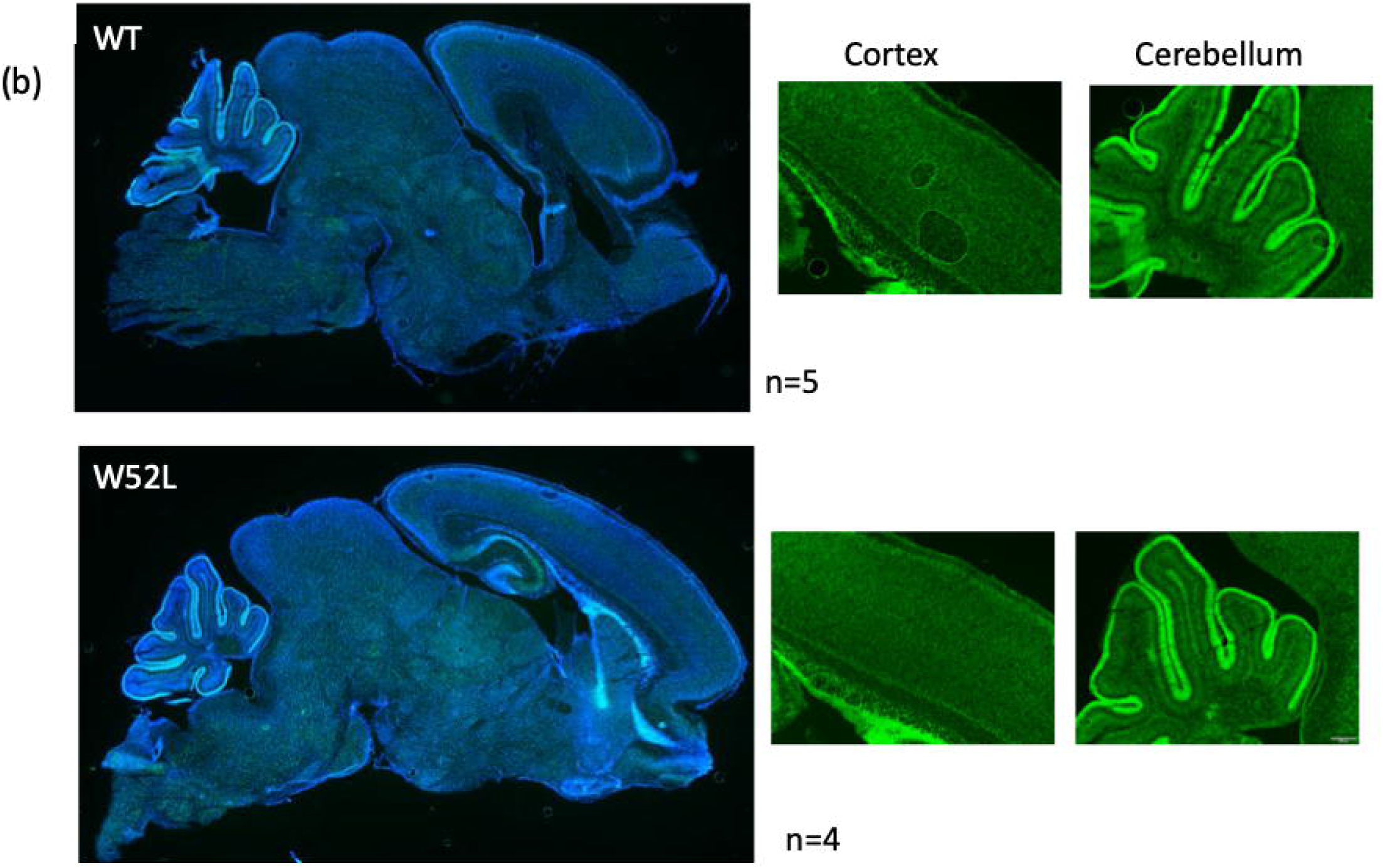
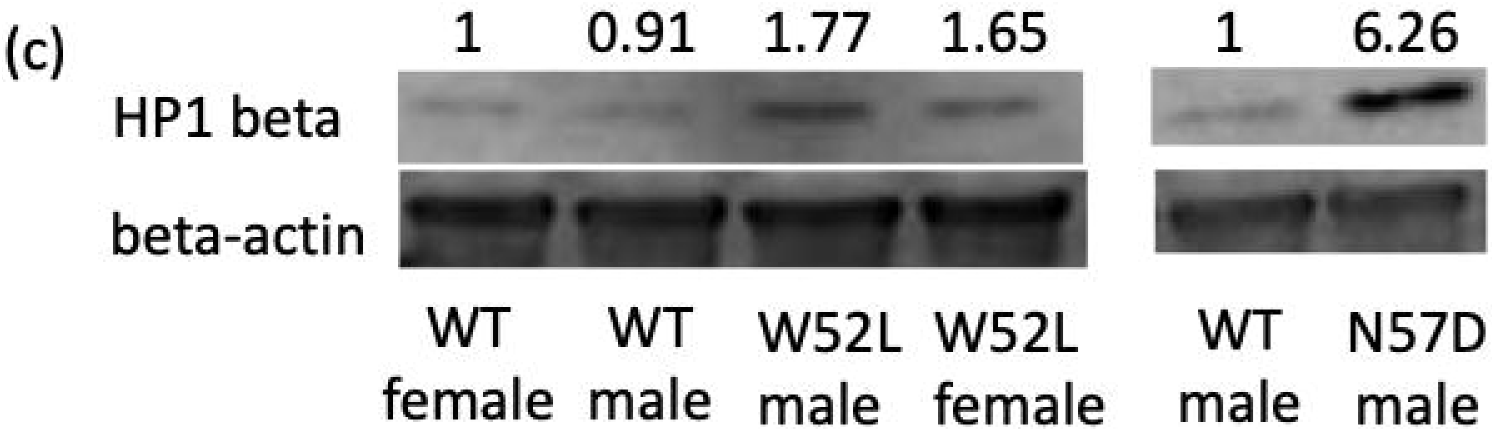
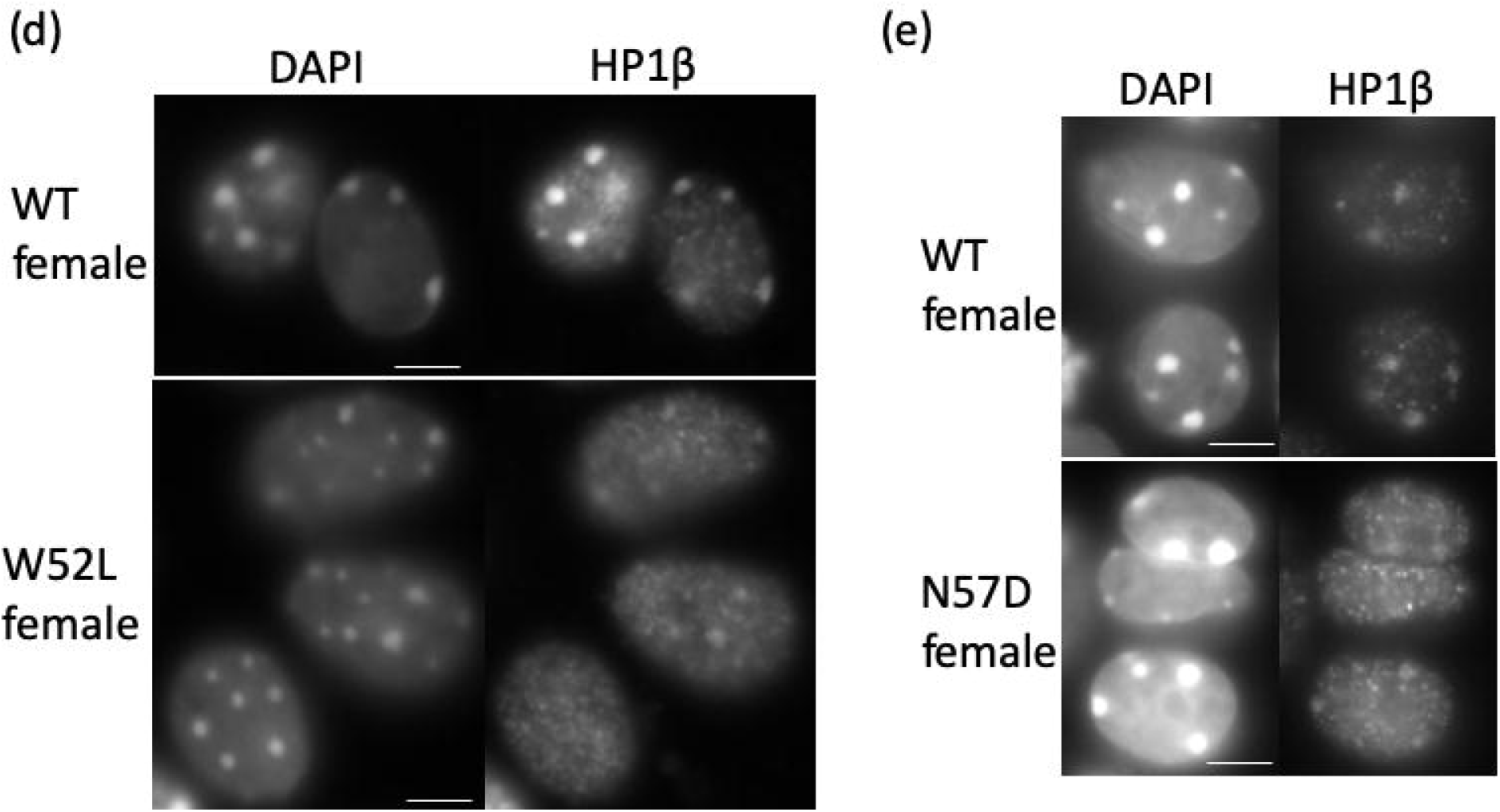
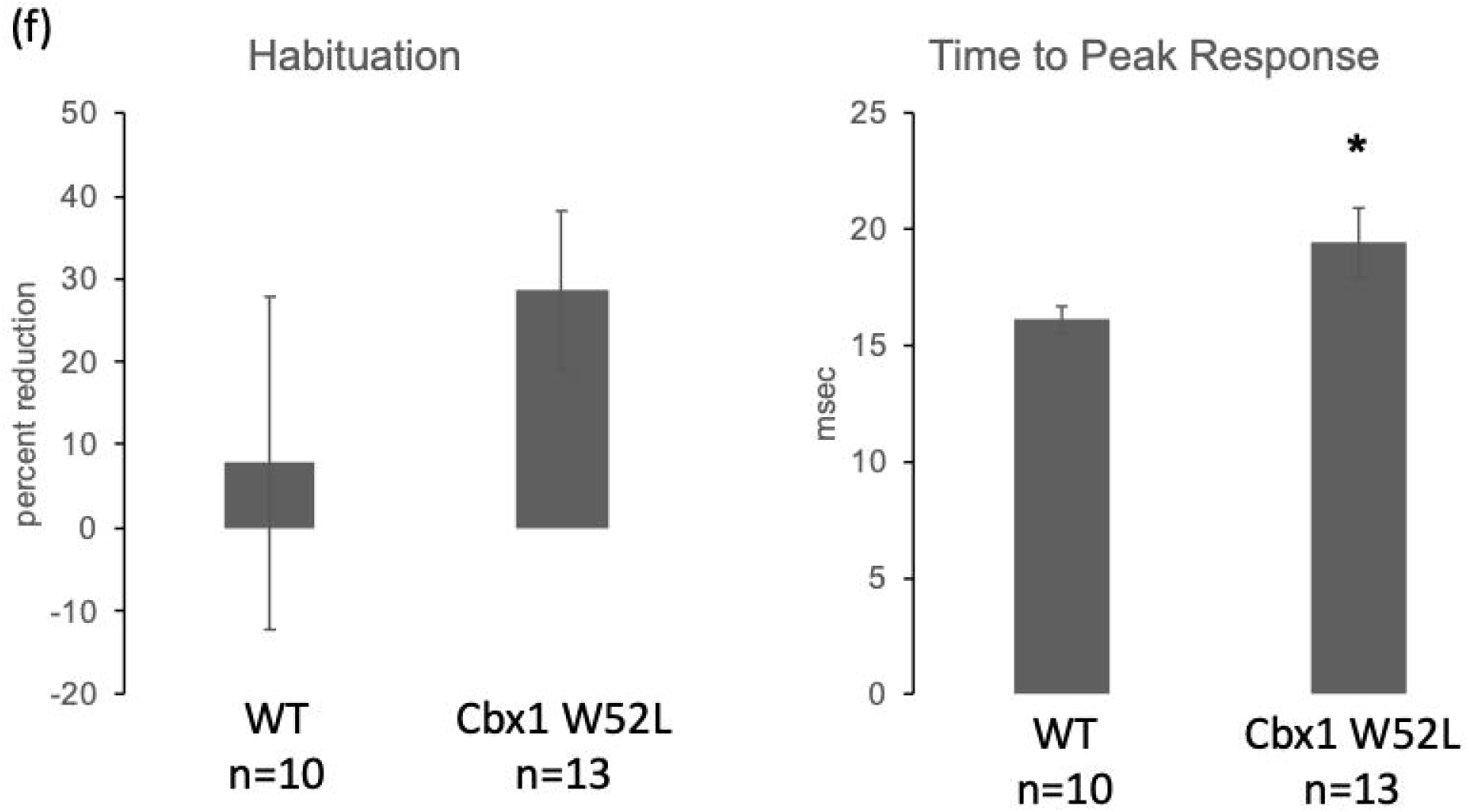
Evaluation of CD *Cbx1* mutant mice. (a) Conservation of amino acid sequences in HP1 β between human and mouse. (b) Nissl staining of *Cbx1* W52L +/− brain at P5 littermates (WT male n=2, WT female n=3, W52L male n=1, W52L female n=3). No obvious morphological differences were noted between WT and W52L mutant tissue samples. (c) Immunoblotting of HP1β using *Cbx1* W52L and N57D +/− P5 neonate forebrain tissue samples. HP1β expression levels were quantified using beta-actin as internal controls. *Cbx1* mutant sample demonstrated higher HP1β protein abundance compared to WT controls. (d) Immunofluorescence staining of HP1β in *Cbx1* W52L +/− primary culture neuronal cells for 12 days *in vitro* (DIV 12). Chromocenters were seen in DAPI-stained nuclei. While HP1β colocalized with chromocenters in neuronal cells obtained from WT mice, HP1β staining of *Cbx1* W52L mutant sample revealed a lack of distinctive colocalization. (e) Immunofluorescence staining of HP1β in *Cbx1* N57D +/− primary culture neuronal cells. Similar results were obtained by using *Cbx1* N57D +/− mutant mice. (f) Behavioral testing of *Cbx1* W52L +/− mutant mice at 4 months. Acoustic Startle Response for Habituation and Latency to Peak Response are shown. A rudimentary assessment of habituation is obtained by comparing the response to 120dB white noise bursts delivered at the beginning and end of the acoustic startle response / pre-pulse inhibition (ASR/PPI) session. The time to the peak response to the first block of 120dB stimuli is also determined. There is no significant difference between wild-type and *Cbx1* W52L +/− mice on habituation to repeated stimuli. However, a significant difference is found in the latency to peak response, p<0.05.

Consistent with the findings in human patients with *CBX1* mutations, the brain structure of *Cbx1* W52L mice at P5 showed no abnormalities compared with WT controls, based on Nissl staining (**Fig 4**). Similar to the findings in patient LCLs, HP1β was increased in the mutant P5 forebrain (cortex and hippocampus) of *Cbx1* W52L and N57D heterozygous mutant mice compared to that of the control mice (**Fig 4c**).

Cultured primary neuronal cells were dissected from the P5 brain of *Cbx1* mice at 7-21 days to evaluate the effects of heterozygous *CBX1* CD variants on HP1β distribution in neuronal cells. In control WT neuronal cells, HP1β colocalized with chromocenters stained with DAPI in the nuclei of cells (**Fig 4d**). In contrast, W52L and N57D heterozygous mutant samples demonstrated that HP1β localized throughout the nucleus without distinct accumulation at chromocenters seen in WT, despite the presence of WT HP1β, suggesting these *Cbx1* mutations exert a dominant-negative effect (**Fig 4d and 4e**).

Behavioral assessments were performed to evaluate the general health and reactivity of our *Cbx1* W52L mouse line. The mutant mice did not differ from their WT littermates in open-field ambulation, rearing, or center activity (**Supplementary Fig 3**). Mutant mice also demonstrated normal balance and coordination in the accelerating rotarod test (**Supplementary Fig 3**). Spontaneous alternations in the Y-maze were did not differ from controls, consistent with shortterm memory was intact in the *Cbx1* mouse mutants (**Supplementary Fig 3**). Finally, an acoustic startle with prepulse inhibition was performed to assess reactivity in the mutant line, which showed typical reactivity and sensory-motor integration. The *Cbx1* mutant mice showed an increased latency to the peak startle response (**Fig 4f**). This slower response is consistent with delayed neurotransmission.

## Discussion

Here, we report identification of *de novo CBX1* missense variants in three individuals with developmental delay, hypotonia, and autism spectrum disorder. Structure/function assays of the identified *CBX1* variants support the notion that these variants disrupt interaction between HP1β and methylated histones. Finally, our novel *Cbx1* mutant mice possessing CD variants demonstrate increased latency-to-peak acoustic response, indicating HP1β function is required for neurofunctional regulation. These data support HP1β having a non-redundant role in brain development, despite the presence of two other closely related homologs (HP1 α and HP1γ).

The HP1β protein consists of an N-terminal chromodomain (CD) (amino acids 21-70) and a C-terminal chromoshadow (CSD) domain (122-168), which facilitates dimerization of HP1 homodimers and heterodimers.^26,27,33–35^ The N-terminal HP1β CD facilitates binding to H3K9me2 or H3K9me3 histone marks^1,36^ via an aromatic cage containing the residues Y21, W42 and F45.^14,37^ The histone binding interaction however also involves other residues in HP1β, including a hydrogen bond between E53 and histone S10, as well as interactions from T51 and W52 that act to stabilize E53. H3K9 methylation is associated with chromatin silencing, and HP1β recognition of this methylated histone mark is necessary for this process. Complementary interactions between HP1β CD and adjacent residues of the H3 tail also play an important role in heterochromatin formation.^14^

It is of note that the *CBX1* mutations identified in this study result in the substitution of amino acids that play key roles in mediating interactions between HP1β and methylated histones. The W52 residue is adjacent to T51, which is phosphorylated by casein kinase II in HP1β, which has a critical effect on nucleosome binding properties.^38^ These properties are important for mobilization of HP1β and lead to phosphorylation of histone H2AX and initiation of the DNA damage response.^39^ Directed evolution testing modelling has previously shown a strong preference for asparagine and alanine at position 57.^40^ The locations of the mutated amino acids identified in this study (T51, W52, and N57) confirm the importance of HP1β-methylated histone interactions in neurocognitive development.

Functional data are consistent with the identified *CBX1* mutations having a dominant-negative mechanism of function. The ChIP-Western blot confirmed that mutant HP1β loses its ability to bind to chromatin, resulting in intranuclear distribution alterations of the mutant HP1β. As the HP1β CD mutants preserves the ability to interact with other HP1 proteins and interacting chromatin proteins, alteration of the intranuclear distribution of HP1β likely affects the location of HP1 interacting proteins, resulting in dominant-negative effects (**Supplementary Fig 4**). The lack of HP1β foci in heterozygous *Cbx1* mutant neuronal cells confirms that mutant HP1β interferes with the proper localization of WT HP1β through a dominant-negative mechanism.

While this dominant-negative mechanism is expected to impact intranuclear localization of most HP1β-interacting proteins, it is worth noting that the HP1β IP-MS experiments showed that the CD mutant HP1β demonstrated reduced GLP/G9a and PRC1.6 complex binding. Both complexes are involved in chromatin organization. GLP/G9a complexes possess lysine methyltransferase activity and are involved in transcriptional regulation.^29–31^ PRC1.6 is a non-canonical polycomb repressive complex 1 that participates in transcriptional repression.^32^ The reduction of GLP/G9a complex binding could be explained by the fact that EHMT1 harbors H3K9m3 mimic sequences that are bound by HP1.^5^ The reduction in PRC1.6 complex binding in CD mutant HP1β suggests the importance of the CD in mediating HP1β and PRC1.6 complex interactions. Therefore, it is possible that chromatin binding of GLP/G9a and PRC1.6 complex could be particularly impaired by the CD mutant HP1β.

Mutated *CBX1* abolishes the interaction of HP1β with GLP/G9a and PRC1.6 complexes, raising the possibility of a shared molecular pathogenesis between several neurodevelopmental disorders. The GLP/G9a complex including EHMT1, is mutated in Kleefstra syndrome^6,7^ and ZNF462, is mutated in Weiss-Kruszka syndrome.^10^ The PRC1.6 complex includes *RNF2*, is mutated in Luo-Schoch-Yamamoto syndrome.^11^ All these syndromes are characterized by developmental delay, intellectual disability, and autistic features, similar to those seen in individuals with *CBX1* mutations. Therefore, disruption of this common molecular pathway may underlie the neurocognitive impairment seen in individuals with *CBX1, EHMT1, ZNF462*, and *RNF2* mutations.

While all three *CBX1* CD mutations resulted in comparable neurodevelopmental phenotypes in humans, proteomics data detected some notable biochemical differences between the N57D and W52L variants. N57D demonstrated a major reduction in HP1β-G9a or PRC1.6 complex binding, while W52L demonstrated milder defects. While the identified pathogenic human *CBX1* variants reside within the CD, it remains possible that variants outside of the CD or loss-of-function variants also result in a human developmental disorder. In support of this possibility, in the gnomAD population database, *CBX1* loss-of-function variants were not identified (11.5 expected, 0 observed, pLI score of 0.98), and the number of observed missense variants was fewer than expected (99.3 expected and 28 observed, Z=2.54).^19^

The establishment of animal models of neurocognitive disorders advances the understanding of their etiology by providing valuable tools for exploring therapeutic interventions. Novel mouse models recapitulating the relevant *Cbx1* mutations, N57D and W52L, were therefore created. Previously, a mouse model with homozygous *Cbx1* loss-of-function multi-exon deletion was shown to have abnormal neocortical columnar organization and were also found to have defective neuromuscular junction development.^12^ The phenotypic difference between heterozygous *Cbx1* CD mutant and loss-of-function mutant mice could be explained by dominant-negative effects. As HP1 forms homo-and heterodimers, any HP1 dimers incorporating the CD mutant HP1β have a diminished ability to bind chromatin (**Supplementary Fig 4**). Hence, the available functional HP1 dimers in HP1β CD mutants are expected to be fewer than those in the heterozygous HP1β knockout model. Despite this mechanistic difference, the latency-to-peak acoustic response was increased, which is consistent with impaired neuromuscular junction function reported in *Cbx1* loss-of-function mutant mice.^12^ These new *Cbx1* mutant mouse lines will serve as relevant models to explore HP1β-chromatin regulation in mammalian neurocognitive development.

In summary, we report identification of dominant-negative *CBX1* missense variants within the HP1β CD in individuals with neurocognitive disorders. This represents the first genetic disorder caused by germline mutations in a gene encoding HP1 proteins. These data are consistent with the important role HP1β plays in neurocognitive development via regulation of heterochromatin formation. Moreover, the locations of the identified *CBX1* mutations indicate the critical importance of HP1β-chromatin binding in precisely regulating global chromatin organization in humans. Further studies to elucidate the molecular pathogenesis of *CBX1*-related syndrome will uncover the significance of HP1-mediated heterochromatin organization in human development.

## Data Availability

Data supporting this work is available upon request. Clinical data sets have been deidentified.

## Acknowledgements

We thank the individuals with *CBX1* variants and their families who participated in this research study. We thank Sho-Hei Noguchi and Akemi Murakami for technical assistance with proteomic analysis. The research was in part supported by funding from the Australian National Health & Medical Research Council to The Centre for Research Excellence in Neurocognitive Disorders (APP1117394). The authors thank the GeneMatcher website for facilitating the collaboration.^13^ Behavior procedures were performed with the assistance of the Neurobehavior Testing Core at UPenn/ITMAT and IDDRC at CHOP/Penn P50HD105354.

## Ethics Declaration

This study was reviewed and approved by the Children’s Hospital of Philadelphia Institutional Review Board (IRB) (#16-013231) for patients identified through the institution. Each patient was enrolled in appropriate local IRB studies per their institutional requirements. Informed consent was obtained from all participants as required by the IRB and local institutional requirements. Clinical data has been deidentified. Written consent for inclusion in this publication was obtained for all participants. This study adhered to the principles set out in the Declaration of Helsinki. Animal experiments were approved by the Institutional Animal Care & Use Committees (IACUC) of the Children’s Hospital of Philadelphia (#21-001291) and the University of Pennsylvania Perelman School of Medicine (#804484).

