## Supplemental Document for "Dominant-negative mutations in *CBX1* cause a neurodevelopmental disorder"

### Supplementary Figures

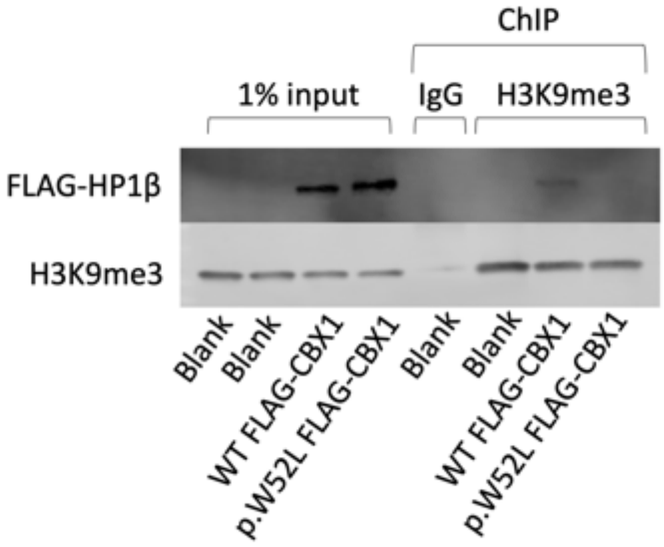


**Supplementary Fig 1:** Reduced p.W52L HP1β binding to H3K9me3 marked histones. Reduced H3K9me3 binding of p.W52L mutant HP1β. ChIP was performed after 48 hours of FLAG-CBX1 cDNA overexpression (wild type and W52L mutant). Although an equal amount of FLAG-CBX1 is expressed between wild-type and W52L mutant, less W52L mutant was identified in the H3K9me3 marked chromatin fraction compared to controls. Biological duplicates revealed a consistent result.

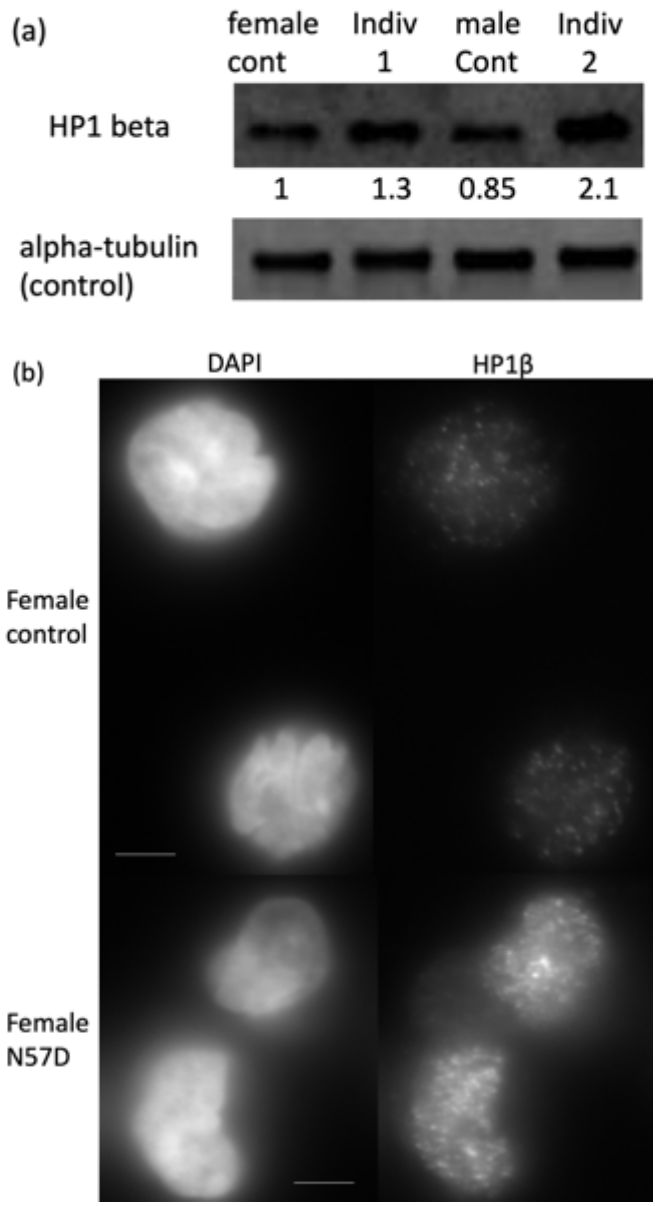


**Supplementary Fig 2:** Accumulation of HP1β in the patient-derived LCLs with *CBX1* mutations. (a) Immunoblotting of HP1β revealed mildly increased HP1β protein in the patient-derived LCLs compared to controls. (b) Representative images of immunostaining of HP1β using control and patient LCLs showing the amount of HP1β is greater in patients than in controls. Biological duplicates revealed a consistent result. Scale bar is 5µm.

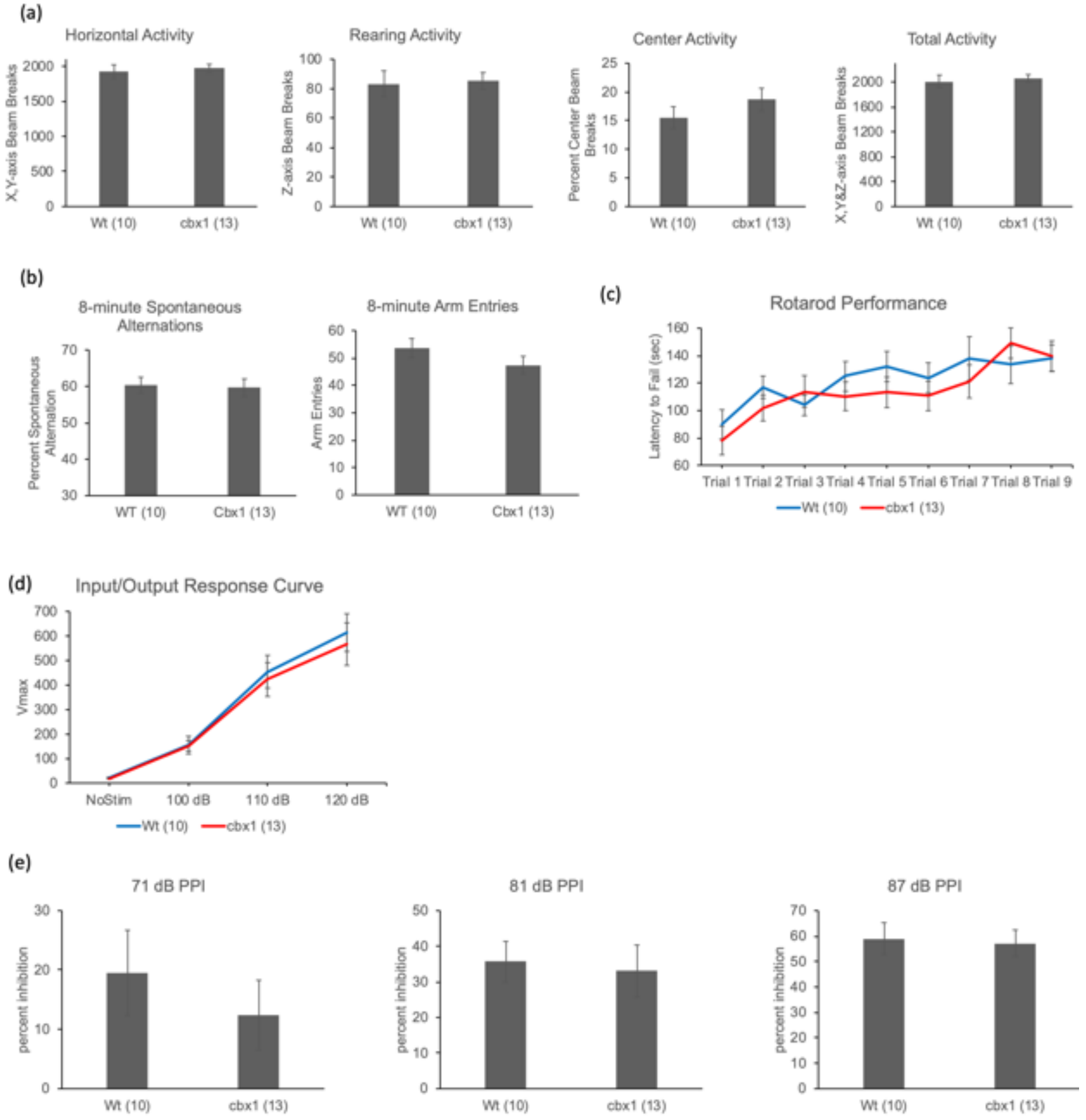


**Supplementary Fig 3**: Behavioral testing results of *Cbx1* W52L+/- mutant mice (at 4 months). (a) Open field activity. Wild-type mice and *Cbx1* W52L +/- mice show typical locomotor and rearing behavior, demonstrating good general health and a propensity to explore the arena. Wild-type mice and *Cbx1* W52L +/- mice show typical percent activity in the center of the arena. *Cbx1* W52L +/- mice showed slightly increased center activity but without statistical significance. The combined horizontal and rearing activity is not different between the genotypes. (b) Y-maze

Spontaneous Alternations (measuring working memory deficits). Both genotypes had similar entries into the maze arms showing similar exploration drive and locomotion (c) Accelerating Rotarod. Wild-type mice show the typical increased latency to fail. The *Cbx1* W52L +/- mice performed similarly to WT so the balance and coordination are normal. In addition, *Cbx1* W52L +/- mice also improve their performance over trials showing normal motor learning ability.

(d) In the stimulus intensity/response curve, Wild-type mice and *Cbx1* W52L +/- mice show the anticipated increased startle response to increased decibels demonstrating that the neural circuitry for response is intact. In pre-pulse inhibition, Wild-type mice and *Cbx1* W52L +/- mice show typical increased inhibition as the intensity of the pre-pulse stimuli is increased demonstrating normal sensory-motor integration.

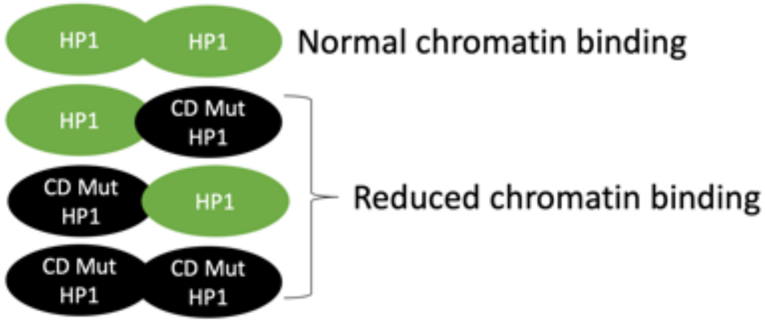


**Supplementary Fig 4:** Model of HP1 dominant-negative effects.

**Supplementary Table 1:** Mass spectrometry analysis of three replicates of Flag-HP1β immunoprecipitates. The scores (IP efficiency and reproducibility) for identification of HP1β-interacting proteins, numbers of unique spectra, and log10-transformed emPAI values for 142 HP1β-interacting proteins are shown.

### Supplemental Methods

#### Exome sequencing:

Individual 1: Exome sequencing was performed as previously described.[1-3] Briefly, genomic DNA was extracted from peripheral blood following standard DNA extraction protocols. Targeted exons were captured using the Agilent SureSelect XT Clinical Research Exome Version 1 kit and sequenced on the Illumina HiSeq 2500 platform with 100 bp paired-end reads. Sequence data was processed using an in-house custom-built bioinformatics pipeline.

Individual 2: DNA was extracted from peripheral blood, and exome sequencing was performed using CRE II whole exome kit (Agilent) on a NovaSeq 6000 instrument (Illumina) with variant annotation, filtering, and prioritisation were performed using an in-house genomics pipeline, Genomics Annotation and Interpretation Application (GAIA) as described previously.[4] Variants were prioritised using an automated PubMed search linked to the referral phenotype performed within GAIA as well as annotation from clinical genetic and variant databases, including ClinVar (http://www.ncbi.nlm.nih.gov/clinvar), HGMD (http://www.hgmd.cf.ac.uk/ac/index.php) and OMIM (http://www.omim.org).

Individuals 3: DNA library underwent exome capture using SureSelect Human all Exon V6 (Agilent). Samples are sequenced on a HiSeq2500 using a 100bp paired-end flowcell (Illumina). Sequencing data analysis was performed using Churchill (PMID:25600152) and variant annotation was performed by VarHouse.

#### Immunoblotting:

Patient LCLs were cultured in RPMI, 20% FBS culture medium at 37°C in 5% $CO_2$. 5.0 × $10^6$ LCL cells were harvested and lysed in 200 µl of fresh, prechilled RIPA buffer (20 mM HEPES pH 7.4, 10 mM KCl, 100 mM NaCl, 1.5 mM $MgCl_2$, 0.34M sucrose, 10% glycerol, 10 mM NaF, 10 mM beta-glycerophosphate, 10 mM sodium butylate, 1 mM DTT, 0.25% Triton-X) with complete protease inhibitor for 10 min on ice. 90 µl of lysate were taken as whole-cell lysate. The remaining samples were centrifuged at 4000 rpm for 5 min at 4°C and the supernatant was transferred to 1.5 ml tubes. Supernatant protein concentrations were measured using the Pierce BCA protein Assay kit (Thermo Fisher Scientific, 23225).

Lysates were mixed with 4× Licor Protein Loading Buffer (928-40004) and incubated at 96°C for 5 min. Protein lysates were then subjected to standard sodium dodecyl sulphate-polyacrylamide gel electrophoresis (SDS-PAGE) using 4-15% polyacrylamide Tris/glycine gels (Bio-Rad, 4561086). Proteins were then blotted to PVDF LF membrane (Bio-Rad, 1620264) using Biorad's Trans-Blot Turbo system (Bio-Rad, 1704150) and blocked for 1 hour in TBS Intercept Blocking Buffer (Licor, 927-60001) and subsequently immunoblotted at 4 degrees overnight using the antibody of choice diluted 1:1000 in blocking buffer. After washing in TBST, membrane were incubated with the IRDye secondary antibody for 1 hr and imaged using the Odyssey imaging system (Licor Model 9120). Antibodies used for immunoblotting were alpha tubulin: Sigma-Aldrich B-5-1-2; HP1 beta, Cell signaling, D2F2; beta actin, Cell signaling 8H10D10. Band intensity quantification were calculated using Licor Image Studio Lite software.

**Chromatin immunoprecipitation (ChIP)-western blot (WB):**

ChIP-WB was performed according to the protocol described previously.[5] Following 48 hours after the CBX1 cDNA vector transfection, cells were fixed with 1% of formaldehyde for 5-10 min. Cross-linking reaction was quenched using 2.5 M glycine. Cells were washed with cold PBS twice, and then PBS or lysis buffer 1 (LB1; 20 mM Tris–HCl, pH 7.5, 10 mM NaCl, 1 mM EDTA, 0.2 % NP-40, 1 mM PMSF) was added. Cells were harvested and were centrifuged at 1,500 g for 5 min at 4°C. The supernatant was discarded, and pellets were washed with 1 mL of PBS and transferred to 1.5 ml tubes followed by centrifugation at 5,000 g for 5 min. For the immunoprecipitation, Protein A (for rabbit antibody) or Protein G (for mouse antibody) magnetic beads were washed with BSA/PBS twice at 4°C. H3K9me3 (Abcam: ab8898), H4K20me3 (Abcam: ab9053) and H3K27me3 (Active Motif: MABI0323) antibodies were added to Protein A/G magnetic beads, and they were rotated at 4°C for >3 hours. After the bead-antibody reaction, they were washed twice by BSA/PBS, and once with LB 3 (20 mM Tris–HC l, pH 7.5, 150 mM NaCl, 1 mM EDTA, 0.5 mM EGTA, 1% Triton X-100, 0.1% Na-Deoxycholate, 0.1% SDS, protease inhibitors), and resuspended in 100 µl of LB3. Collected cells were resuspended with 1 ml of LB1 and were lysed on ice for 10 min. After centrifugation at 2,000 g for 5 min, pellets were resuspended in 1 ml of LB2 (20 mM Tris–HCl, pH 8.0, 200 mM NaCl, 1 mM EDTA, 0.5 mM EGTA, 1 mM PMSF). The tubes were placed on ice for 10 min, then spun again to remove supernatant. Obtained pellets were lysed with 1 ml of LB3 for 10 min. After the reaction, the tubes were centrifuged at 2,000 g for 5 min, and the pellet was resuspended in 400 µl of LB3, and placed on ice for 10 min. After centrifugation at 2,000 g for 1 min, obtained pellets were sonicated and centrifuged at 20,000 g for 15 min and supernatant was transferred to new tubes, 30 µl aliquots were taken as whole cell lysate (WCE) samples and additional 10 µl aliquots for Western Blot. Harvested input samples were mixed with antibody beads complexes and were rotated overnight at 4°C. Magnet beads were collected using magnetic stands and washed with 1 mL cold RIPA buffer five times and then washed once with 1 mL cold TE50. After this, the beads were centrifuged at 2,000 g for 1 min, and placed in a magnetic holder. Harvested washed magnet beads were resuspended in 50 µl of elution buffer. Then, 2× Laemmli sample buffer was added to the beads, and they were placed in a 95°C heat block for 30 min with vigorous vortexing every 5 min. Tubes were then centrifuged at 2,000 g for 1 min, and the soluble ChIP lysates were transferred to a new tube. Collected ChIP samples were used for Western blotting.

**Immunofluorescence (IF):**

IF was performed as previously described.[6] HEK293T cells were settled on poly-lysine coated coverslips and transfected with the CBX1 expression vector. Cells were fixed in 4% formaldehyde in PBS for 10 min at room temperature (RT) 48 hours after the transfection, permeabilized in 0.5% Triton X-100 in PBS for 5 min at RT, then incubated with IF block (2% FBS, 2% BSA, 0.1% Tween,

0.02% Sodium Azide in PBS) for 20 min at RT. Cells were incubated with a mouse primary antibody against FLAG (Sigma, F1804) for 1 hour at RT, followed by three washes in 0.2% Tween in PBS. Subsequently, cells were incubated with Alexa Fluor 488 donkey anti-mouse IgG (1:500) for 1 hour at RT. After three washes in 0.2% Tween in PBS, and one wash with PBS, cells were mounted in Vectashield containing 4′,6-diamino-2-phenilinodole (DAPI, Vector Laboratories, H-1200). Images were captured on a Leica wide-field fluorescence microscope, using a 1.4 NA 63× oil-immersion objective (Leica) and an ORCA-Flash4.0 V2 Digital CMOS camera (Hamamatsu, C11440-22CU), controlled by LAS X software (Leica).

**Proteomics analyses:**

To establish stable cell lines for inducible expression of Flag-tagged HP1β, the Flp-In T-REx 293 system (Invitrogen) was used as described.[7] Protein extraction, immunoprecipitation, and mass spectrometry were performed as described.[7] Briefly, a cell extract was prepared from ~1× $10^8$ cells using CSK buffer containing 0.5M NaCl and incubated with 40 µl anti-Flag agarose beads (Sigma). The bound proteins were eluted by incubating with 0.2 mg/ml 3×Flag peptide (Sigma) and separated in a 12.5% SDS-polyacrylamide gel for liquid chromatography coupled to tandem MS (LC/MS/MS). Data analysis for LC/MS/MS was performed as described previously, with minor modifications.[7] The International Protein Index database (Human, version 3.87, ftp://ftp.ebi.ac.uk/pub/databases/IPI) was used. The number of unique spectra for each protein were counted from data summarizing two measurements from all pieces of a gel and converted to emPAI (exponentially modified protein abundance index) value [8]. HP1β-interacting proteins were identified using two scores, IP efficiency and reproducibility, according to identification of HP1α-interacting proteins.[7] For calculation of IP efficiency, only immunoprecipitated samples of HP1β WT and V23M were considered. Using the same threshold as for HP1α and excluding HP1 and histones, we identified 142 HP1β-interacting proteins (Supplemental table 1). For Fig 4a, the fraction of HP1 was calculated as the number of unique spectra specific for the HP1 subtype divided by the total number of unique spectra for all HP1. To classify HP1β mutants in Fig. 4d and e, log10-transformed emPAI values of 142 HP1β-interacting proteins were used as a profile. For Fig. 4f, IP enrichment of each protein was defined as the log10-transformed ratio of the average emPAI value between three replicates of the Flag-HP1β immunoprecipitates to that of the mock immunoprecipitates.

**Animal ethics:**

All experiments were performed using the C57BL/6N strain obtained from Charles River. Wild type and genetically engineered *Cbx1* variants were maintained on a 12 h light / dark schedule. Mice were bred and maintained in the animal care facilities of the Children's Hospital of Philadelphia and the University of Pennsylvania. All animal procedures and experiments were approved by the Institutional Animal Care and Use Committees of the Children's Hospital of Philadelphia or the University of Pennsylvania.

**Generation of *Cbx1* mutant mice by CRISPR/Cas9 genome editing:**

CRISPR/Cas9-mediated gene editing was conducted in zygotes of C57BL/6 mice. A guide RNA targeting a genomic region of interest, Cas9 and ssDNA containing mutant sequence were electroporated into zygotes. CrisprRNAs and tracr RNAs from Integrated DNA Technologies were used. For W52L mutagenesis, the following gRNA and ssDNA sequences were used. gRNA: GTTTCAGTGAGGACAACACT and ssDNA: TGATCTGAATGGAAAGTATAACTTGCTGTGCTTGCTGGTTTCAGTGAGGACAACACTTTGG AGCCAGAAGAGAATCTGGATTGCCCTGACCTTATTGCTGAGTTTCTACAGTCACAGAAA. As the single nucleotide variant (SNV) we intended to introduce span within the PAM sequence, we did not introduce additional SNV within the template ssDNA. For N57D mutagenesis, the following gRNA and ssDNA sequences were used. gRNA: TTGGGAGCCAGAAGAGAATC, and ssDNA: GTATAACTTGCTGTGCTTGCTGGTTTCAGTGAGGACAACACTTGGGAGCCAGAAGAGGATC

TAGATTGCCCTGACCTTATTGCTGAGTTTCTACAGTCACAGAAAACAGCTCATGAGACA. To prevent re-cut of the target site, an additional silent SNV was included in the ssDNA. Mice were screened for the introduction of the desired mutation by PCR. Sanger sequencing of PCR products was used to confirm genomic incorporation of the desired mutation. Founder mice were backcrossed to C57BL/6 wild type mice for at least two generations before being used in experiments.

**Primary neuronal culture:**

P5 littermate mice were euthanized by decapitation. Whole brain was dissected, cut into small pieces, and collected into prechilled HBSS in 15 ml tubes. Tissues were dissociated in 0.25% Trypsin-EDTA and 5 μg/ml DNase I at 37 degrees for 20 min, and then gently triturated. After filtering using cell strainer and washing several times with DMEM / F12 / 10% FBS, cells were spread on a poly-D-lysine coated cover glass and cultured for 7-22 days in growth medium (DMEM / F12 / 2% of B27 supplement).

**Nissl staining:**

Nissl staining was performed as previously described.[9] Briefly, 20-μm-thick was prepared from the cryopreserved brain tissue, and sections were stained with cresyl violet (MP Biomedicals).

**Open-field activity:**

Spontaneous activity in an open-field arena is used to assess ambulation and rearing as part of a general health evaluation. After 30 minutes of habituation to the procedure room, a mouse is placed in the center of the Plexiglas arena (14 in. x 14 in. x 18 in) for a 10 min trial. IR emitters and detectors collect beam breaks as activity data to record rearing, center and peripheral activity (Photobeam Activity System, San Diego Instruments).

**Y-maze task:**

After habituation to the procedure room, an individual mouse is placed at the distal end of an arm in a maze with three arms in a Y shape. The mouse explores the maze for eight minutes with each arm noted (all four paws are in that arm), When exploring, mice have a natural propensity to avoid the arm just vacated and enter the other arm that is they alternate their entry into the arms of the maze. A reduction is spontaneous alternations is considered an impairment if short term/working memory. Percent spontaneous alternations is calculated as % Spontaneous Alternation= [ Number of alternations / (Total arm entries−2)] ×100.

**Accelerating rotarod:**

The Rotarod (IITC San Diego Ca.) is a 1-inch diameter, horizontal rod with a rough surface that set to rotate and/or accelerate at different rates. As the rod speed increases, the mouse must adjust its stride to remain walking on the rod. An inability to adjust its cadence will cause the mouse to fall or grip onto the rotating rod. A decrease in the latency to stop walking on the Rotarod suggests a coordination or balance impairment. Mice receive three trials per day over three consecutive days with the Rotarod programed to accelerate from 4 to 40 rpm during a 5 minute trial. Note that on the first day only, mice were place on the stationary rod to allow for habituation before rotation starts. All intertrial intervals were about 30 minutes. The equipment was wiped with 70% EtOH between all trials. Every test day, mice were habituated to the procedure room for thirty minutes. Latency to fail was defined as the time to drop from the rod or time to make a full rotation while gripping on to the rod.

**Acoustic startle response/prepulse inhibition (PPI) and habituation:**

The acoustic startle response (ASR) is a standard test of motor responsiveness to strong sensory stimuli. Pre-pulse inhibition (PPI) is included in the procedure. PPI naturally occurs when a weaker, non-startle evoking stimulus precedes a stronger startle-evoking stimulus. The weak pre pulse predicts the higher intensity stimuli to attenuate the response and is considered a measure of sensory-motor gating. Deficits in ASR and PPI are seen in post-traumatic stress disorder and schizophrenia. A rudimentary assessment of habituation is also obtained by comparing responses late in the train of stimuli delivered to responses early in the procedure. Mice were tested for ASR and PPI with 4 SR-Lab systems chambers (San Diego Instruments). The chambers are sound attenuating, ventilated boxes illuminated with a 15W light bulb. A 5-inch x 1.75-inch diameter Plexiglas tube is mounted on a platform with a stabilimeter. The stabilimeter transduces motor activity, which is digitized and recorded. Acoustic stimuli are delivered through small speakers mounted inside each chamber. A continuous 70-dB white noise is presented throughout the session. After a 5 min acclimation to the background noise, the mice received a string of 6, 120dB white noise bursts to collect baseline response. This was immediately followed by 40 presentations of 4 different white noise intensities (100,110,120 dB and background noise) spaced about 15 sec. apart. The white noise bursts have 40 msec durations. Each of the intensities are present 6 times in the session in a pseudo-randomized fashion. The movement of the mouse in response to the acoustic stimulus is transduced to a digital format stored on a computer. Activity is sampled for 120 msec beginning at the initiation of the acoustic stimuli. Data is collected as peak amplitude of the startle response, average startle response, total movement during the trial and latency to peak startle response. The data are used to calculate an intensity-response curve. The pre-pulse inhibition (PPI) block of stimuli begins immediately after startle testing is completed. The PPI session consists of 40 120 dB, 40msec white noise bursts spaced 15 sec. apart. The 120 dB stimuli are delivered either alone or preceded by a 20 msec. Pre-pulse stimuli delivered at 3 different intensities (78, 81 or 85 dB). The percent decrease in the startle response with the pre-pulse compared to the startle response without pre-pulse is a measure of PPI. Finally, a string of 6, 120dB white noise bursts are presented to be compared to the block of 120 dB bursts that initiate the session. The latter block in compared to the former to calculate habituation.
